# Resting shear elastic modulus as a marker of peripheral fatigue during maximal isometric contractions in humans

**DOI:** 10.1101/402644

**Authors:** Julien Siracusa, Keyne Charlot, Alexandra Malgoyre, Sébastien Conort, Pierre-Emmanuel Tardo-Dino, Cyprien Bourrilhon, Sebastian Garcia-Vicencio

**Author notes:** **Corresponding authors:** (JS) (SGV).

## Abstract

The aim of this study was to investigate whether the resting *Vastus Lateralis* (VL) muscle shear elastic modulus (µ), evaluated by shear wave elastography, represents peripheral fatigue during repetition of isometric maximal voluntary contractions (MVCs) of the knee extensor (KE) muscles.

Eight healthy well-trained males repeated 60 isometric MVCs of the KE muscles (6 × 10 MVCs; 5 s on/5 s off). Single and double electrical stimulations were delivered to the femoral nerve every ten MVCs during contraction and at rest. The amplitude and properties of the potentiated torque following single (Tw_pot_) double electrostimulation and the amplitude of the concomitant VL compound action potential were considered to be indicators of peripheral fatigue. The resting VLµ was measured during a 5-s rest period after each MVC and electrical stimulation series.

The resting VLµ significantly decreased (-21.8 ± 3.9%; P < 0.001) by the end of the fatigue protocol, decreasing from the 10^th^ MVC to the end of the exercise (60^th^ MVC) for all participants, with the loss ranging from 18 to 29%. The potentiated doublet and single twitch torque (Tw_pot_) decreased by 42.5 ± 10.8% and 55.7 ± 8.8%, respectively, by the end of exercise (P < 0.001 for both). The relative mechanical properties of Tw_pot_, *i.e.* electromechanical delay (P <0 .001), contraction time (P = 0.004), and maximal rate of torque development/relaxation (P < 0.001) also changed significantly during exercise.

This study shows that the kinetics of the resting VLµ is associated with changes in both voluntary and electrostimulated torque amplitudes and electromechanical properties of the single twitch during the repetition of maximal voluntary fatiguing exercise. Changes in the resting VLµ may reflect a decline in muscle function, *e.g*. impairment of excitation-contraction coupling, contractile processes, and/or elastic properties, throughout the increase in muscle compliance, directly affecting force transmission.

## Introduction

Neuromuscular fatigue (NMF) is classically defined as “an exercise-induced reduction in the ability of skeletal muscle to produce power or force, irrespective of task completion” [1]. The study of NMF and underlying mechanisms of recovery, both central and peripheral, is important to prevent overuse injuries [2] and develop new strategies to enhance muscle recovery [3].

NMF and, more specifically, the peripheral modifications that occur within the muscle, at or distal to the neuromuscular junction [4], have been evaluated by artificial stimulation of the skeletal muscle and/or motor nerve structures [5-7]. Modifications in mechanical and/or electromyographic (EMG) response amplitudes delivered by single, double, or tetanic stimulation to a relaxed muscle are considered to be good indicators of peripheral fatigue [1]. More specifically, the study of these electrically-induced responses allows non-invasive investigation of changes of the (i) neuromuscular propagation/transmission of action potentials along the sarcolemma, (ii) excitation-contraction coupling, (iii) cross-bridge cycle, and (iv) intrinsic force [1, 4, 8-11]. Moreover, electromechanical properties of single twitches, such as electromechanical delay (EMD), contraction time (CT), half-relaxation time (HRT), and their first derivates may also reflect mechanical and electrochemical alterations, such as elastic properties, of skeletal muscle related to the capacity of force transmission [12-14].

Aside from contractile mechanisms, the decrease in voluntary and/or electrically induced force with fatigue also depends on the elongation capacity of elastic components, both in series and in parallel, (*i.e*. connective tissue, tendons), in transmitting force [12]. It has been well demonstrated that fatigue can increase muscle-tendon complex compliance due to repetitive cycles of long-lasting contractions [2, 15, 16]. Higher muscle-tendon compliance would furthermore suggest a longer time for force transmission to the bone due to the inability to store and release elastic energy, which can lead to a higher risk of muscle-tendon complex injury [16, 17]. The use of real time ultrasonography has made it possible to assess the displacement of human tendons and aponeurotic structures during voluntary contractions to determine a muscular stiffness index *in vivo* [18-20]. However, these techniques reflect modifications in the stiffness of several structures (muscles, tendons, nerves, and skin) acting around a given joint and not solely stiffness of the muscular tissue. Moreover, conventional ultrasonography techniques often require maximal voluntary contractions (MVC), which may limit its use in clinical cases or acute fatigue states. Thus, the relationship between changes in specific muscle stiffness during fatiguing conditions and its role in muscle performance is therefore unclear and needs to be investigated.

Shear-wave elastography (SWE) is a very promising alternative which provides reliable and quantitative real-time assessment of specific muscular tissue stiffness at rest and during isometric contractions or passive stretching [21-23]. Unlike conventional ultrasound elastographic methods, SWE is not based on manual compression or the extent of tissue displacement [24]. This technique provides a tissue stiffness index based on tissue shear-wave propagation velocity measurements induced by an acoustic radiation force. The shear-wave propagation velocity directly correlates with the muscle shear elastic modulus (µ) in a homogeneously elastic medium, which is assumed to be the case for muscle tissue [25, 26]. Although changes in µ have been well studied after damaging exercise [27-29], there is no consensus concerning its modification after fatiguing conditions. For example, Bouillard *et al.* [30] reported that the dynamic µ, evaluated during voluntary contractions by SWE, closely followed declines in voluntary torque during fatiguing caused by submaximal sustained isometric contractions. Moreover, they showed that its amplitude was affected if fatigue was preliminarily induced *versus* a control non-fatigued session [31]. Andonian *et al.* [32] found that the resting µ of the quadriceps was lower immediately after an extreme mountain ultra-marathon, in which a high level of fatigue is experienced. In contrast, Lacourpaille *et al.* [29] demonstrated that the resting µ was not modified after fatiguing concentric contractions, suggesting that it is not influenced by metabolic changes within the muscle or by peripheral fatigue. However, in their study, 30 isokinetic (at 120°*s^-1^) concentric contractions of the elbow flexor muscles did not induce a significant decrease in voluntary torque, limiting conclusions concerning the relationship between fatigue and changes in resting µ (muscle stiffness). Thus, the kinetics of the resting µ during repeated maximal fatiguing contractions and its relationship with modifications of neuromuscular indicators of peripheral fatigue (*i.e.* changes in mechanical and EMG responses) is still unknown.

The decline in muscle function during and after fatiguing contractions, closely associated with greater muscle-tendon complex compliance [16] and the impairments in mechanical and electrochemical properties of skeletal muscle related to force transmission [12, 13], suggest that repetition of isometric MVCs could induce a progressive decline in resting µ over the fatigue protocol. Thus, modifications of resting µ may be associated with changes in electrically stimulated mechanical (single and double) and EMG (M-wave) responses, which are strongly affected when the muscle is exercised isometrically at maximal intensity for a short time [33, 34]. The aim of this study was to investigate whether the resting µ evaluated by SWE is a good indicator of peripheral fatigue and reflects changes in the mechanical and elastic properties of muscles (*i.e*. increases in muscle compliance) during the repetition of isometric MVCs of KE muscles in humans.

## Materials and methods

### Participants

Eight strength-and-endurance trained French Army soldiers volunteered to participate in the present study. Their anthropometrical and physiological characteristics were as follows: age: 29 ± 2.6 years, weight: 74 ± 5.2 kg, height: 1.7 ± 0.1 m, BMI: 28.8 ± 2.3 kg*m^-2^, VO_2max_ relative to BMI: 53.6 ± 4.7 ml*min^-1^*kg^-1^, and body fat percentage: 11.1 ± 2.9%. They performed regular physical activity, such as strength training, running, and/or cross training (between 6 and 15 h/week), with no recent history of muscular, joint, or bone disorders or receiving any medication that could interfere with neuromuscular responses. This study was part of a medico-physiological follow-up performed at the request of the French Army. All the volunteers were fully informed of the experimental procedures, aims, and risks and gave their written assent before any testing was conducted. Each participant participated in an inclusion session consisting of (i) a complete medical examination with anthropometric data collection, (ii) a standardized incremental maximal aerobic test, and (iii) complete familiarization with the experimental procedures. This study was approved by the scientific leadership of the French Armed Forces Biomedical Research Institute. All experiments were conducted in accordance with the Helsinki Declaration [35].

### Anthropometrical measurements

Body mass was measured to the nearest 0.1 kg using a calibrated scale and height was determined to the nearest 0.01 m using a standing stadiometer. Height and body mass were measured without shoes and while wearing underwear. The body mass index was computed by dividing the mass by the height squares (kg*m^-1^). The percentage of body fat was estimated using skinfold thickness values and Durnin & Womersley standard equations [36]. Skinfold thickness values were measured to the nearest millimeter, in triplicate, at the biceps, triceps, and subscapular, and suprailiac points on the right side of the body using a Harpenden skinfold caliper (British Indicators, West Sussex, UK). The same investigator performed all measurements.

### Intermittent voluntary fatigue protocol

After a 10-min warm-up at submaximal intensity, participants performed an intermittent voluntary fatigue protocol consisting of 6 × 10 repetitions of alternating isometric 5-s MVCs of the KE muscles and 5-s passive recovery periods **(Fig 1)**. The number of contractions was chosen to generate a high level of voluntary strength loss and peripheral fatigue, as demonstrated previously in adults [34]. Double and single electrical stimulations were delivered to the femoral nerve before and every ten MVCs, during the contractions and at rest, to determine the kinetics of central and peripheral neuromuscular fatigue (*see below*). Moreover, a linear transducer was fixed to the skin, using a custom-made system, over the *Vastus Lateralis* muscle (VL) and used in SWE mode (musculoskeletal preset) to capture the shear-wave propagation velocity. After each MVC and electrical-stimulation series, the resting *Vastus Lateralis* shear elastic modulus (VLµ) was measured during a 5-s period to ensure reproducibility of the measurements, as proposed by Andonian *et al.*, [32]. Participants were asked to stay as relaxed as possible and, after stabilization of the elastography 2D color map, a 5-s clip was recorded. The passive right leg KE torque and VL and b*iceps femoris* (BF) EMG activity were evaluated to control for any unwanted muscle contraction in real time. Participants were not informed of the criterion of task failure (60-MVCs) but had visual feedback of the torque output during the exercise. They were also strongly encouraged by the experimenters during the entire fatiguing task.

**Fig 1.**
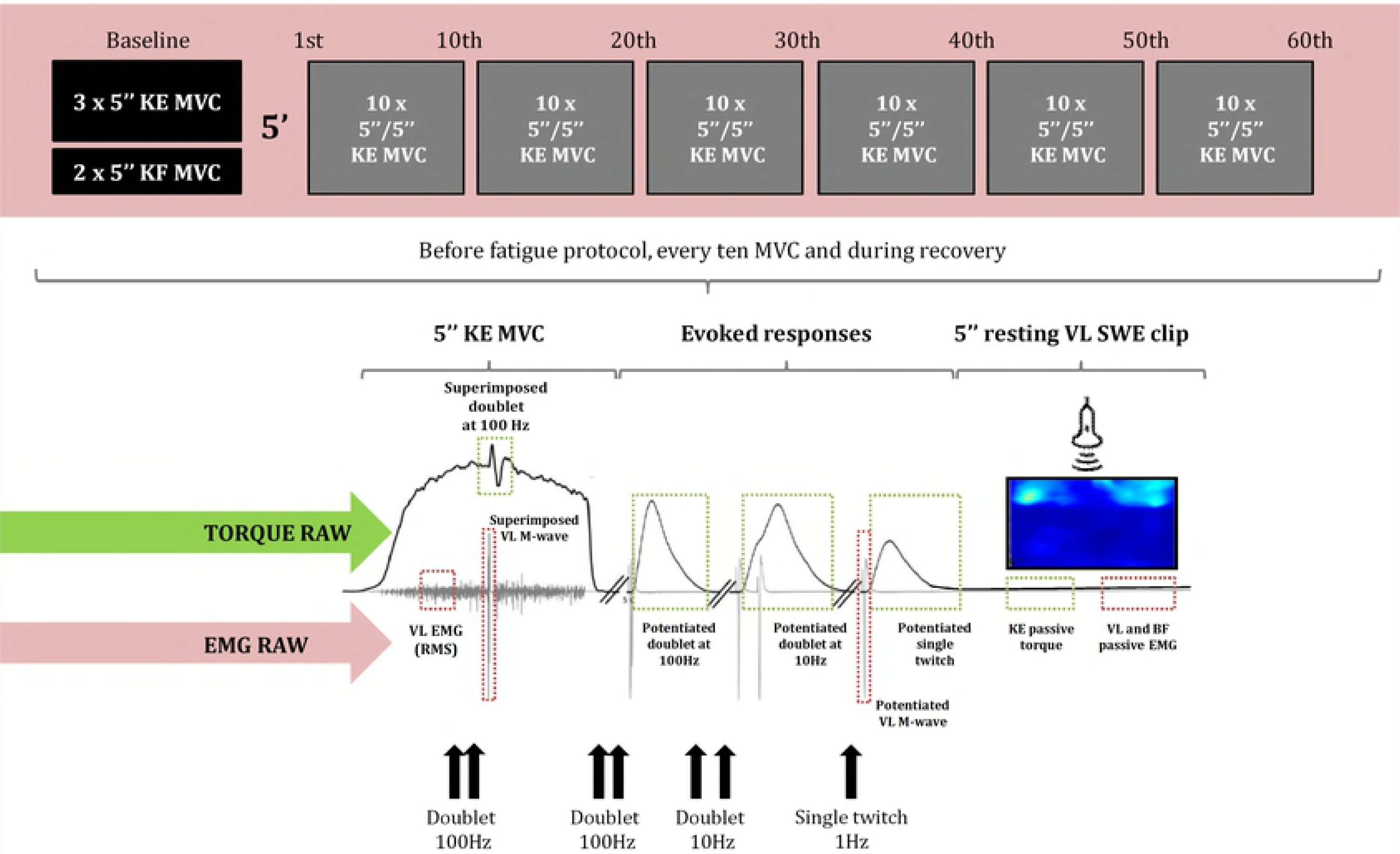
Design of the voluntary intermittent fatigue protocol. This protocol consisted of a series of voluntary force and electrical stimulation and muscle shear-wave elastography measurements performed before the series and during and after every 10 MVC. KE: knee extensors; KF: knee flexors; MVC: maximal voluntary contraction; SWE: shear-wave elastography; EMG: surface electromyography; RMS: root mean square: VL: *Vastus Lateralis*; BF: biceps femoris.

## Neuromuscular function measurements

### Femoral nerve electrical stimulation

Evoked contractions of the KE muscles were triggered with a constant-current stimulator (Digitimer DS7A, Hertfordshire, UK). Single and double square-wave pulses of 1,000 μs, at maximal voltage (400 V), were delivered percutaneously to the femoral nerve using a self-adhesive electrode (10-mm diameter, Ag-AgCl, Type 0601000402, Controle Graphique Medical, Brie-Comte-Robert, France). The cathode was placed in the femoral triangle, 3-5 cm below the inguinal ligament. The anode, a 5×10 cm self-adhesive stimulation electrode (Medicompex SA, Ecublens, Switzerland) was placed at the gluteal fold. Small spatial adjustments were initially performed using a ball probe cathode pressed into the femoral triangle to determine the optimal stimulation site. This site corresponded to the position in which the greatest un-potentiated KE single twitch (T_w_) amplitude and concomitant VL compound muscle action potential (M_max_) amplitude were induced. The optimal stimulation intensity, *i.e.* the intensity at which maximal un-potentiated T_w_ and concomitant VL M-wave amplitudes started to plateau, was determined from a progressive recruitment curve. Briefly, simple pulses were induced every 15 s from 40 to 99 mA in 5-mA increments. The supramaximal stimulation intensity ranged from 52 to 91 mA and corresponded to 130% of the optimal intensity.

### Isometric maximal voluntary contraction

Maximal voluntary and electrically stimulated contractions were assessed under isometric conditions with an isokinetic dynamometer (Cybex Norm, Lumex, Ronkonkoma, NY, USA). Participants were comfortably positioned on an adjustable chair with the hip joint flexed at 30° (0° = neutral position). The dynamometer lever arm was attached 1-2 cm above the lateral malleolus with a Velcro strap. The lever arm was home-built and included a high-density foam pad, placed against the posterior aspect of the leg, and a Velcro strap positioned over the anterior aspect of the leg. This configuration was chosen to reduce cushioning and improve torque transmission and resolution, which is critical when evaluating twitch contractile properties and the voluntary activation level [1]. The axis of rotation of the dynamometer was aligned with the lateral femoral condyle of the femur. The participants were secured firmly with Velcro straps across the hips and chest to minimize upper body movement. During each MVC, participants were instructed to grip the seat to stabilize the pelvis. Before the voluntary fatigue protocol, three 5-s MVCs of the KE and two of the knee flexors (KF) were performed, with 60-s passive recovery periods. The means were defined as the control “non-fatigued” values. Absolute KE and KF torque was determined as the peak force reached during maximal efforts. All measurements were taken from the participant’s dominant leg (right leg for all participants), which was fixed at 90° (0°= knee fully extended). This muscle length was selected to ensure a good reliability of the neuromuscular assessments. Indeed, it is a length close to the optimal angle of force generating capacity and, at this position the quadriceps muscle is relatively lengthened. These last arguments allow us to induce a greater extent of peripheral fatigue and to detect easily muscle stiffness modifications.

Torque data was corrected for gravity using Cybex software and was acquired and digitized on-line at a rate of 2 kHz by an A/D converter (Powerlab 8/35, ADInstruments, New South Walles, Australia) driven by Labchart 8.0 Pro software (ADInstruments).

### Electrically evoked torque and maximal voluntary activation Level

The double pulse (stimulated at 100 Hz) superimposition technique, based on the interpolated-twitch method [37], enabled us to estimate the maximal KE voluntary activation level (VAL). Briefly, superimposed (Db_s100Hz_) and potentiated (Db_pot100Hz_) double stimulations were delivered during MVC after the torque had reached a plateau and 3-s after cessation of the contraction, respectively. This allowed us to obtain a potentiated mechanical response and hence reduce the variability of the VAL values [38]. The superimposed doublet was preferred to a superimposed single T_w_ because it results in a greater signal-to-noise ratio and thus allows the detection of small changes in VAL [39]. The ratio of the amplitude of the Db_s100Hz_ over that of the Db_pot100Hz_ for the relaxed muscle (control doublet) was then calculated to obtain the VAL, (i.e. indicator of central fatigue), as follows:

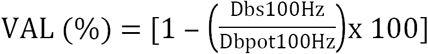

After cessation of the contraction and 3-s after the Db_pot100Hz_, a low frequency (10 Hz) doublet stimulus and a single T_w_ were delivered to the relaxed muscle in a potentiated state (T_wpot_; 3-s between; **Fig 1**) [40]. This set of measurements (MVC with superimposed doublet + evoked stimuli to the relaxed muscle) was repeated before the fatigue protocol and every ten MVCs. Peak torque and the Db_pot10Hz_-to-Db_pot100Hz_ ratio (Db10:100) was then calculated from double pulses; any decrease in this ratio is commonly interpreted as an index of low-frequency fatigue, *i.e.* the preferential loss of force at low frequencies of electrical stimulation [41]. The following parameters were also obtained from the T_wpot_ response: peak torque (Pt), EMD, CT, HRT, maximal rate of torque development (MRTD), *i.e.* the maximal value of the first derivative of the mechanical signal divided by the peak torque, maximal rate of torque relaxation (MRTR), *i.e.* the maximal value of the first derivative of the mechanical signal divided by the peak torque.

### EMG activity

The EMG signals of the VL and BF muscles were recorded, during voluntary and evoked contractions, using bipolar silver chloride surface electrodes (Blue Sensor N-00-S, Ambu, Denmark). The recording electrodes were taped lengthwise to the skin over the muscle belly, as recommended by SENIAM [42], with an inter-electrode distance of 20 mm. The reference electrode was attached to the patella. Low impedance (Z < 5 kΩ) at the skin-electrode surface was obtained by shaving, gently abrading the skin with thin sand paper, and cleaning with alcohol. EMG signals were amplified (Dual Bio Amp ML 135, ADInstruments, Australia) with a bandwidth frequency ranging from 10 to 500 Hz (common mode rejection ratio > 85 dB, gain = 1,000) and simultaneously digitized together with the torque signals. The sampling frequency was 2 kHz. During the fatigue protocol, the root mean square (RMS) values of the VL EMG activity were calculated during the MVC trials over a 0.5-s period after the torque had reached a plateau and before the superimposed stimulation was evoked. This RMS value was then normalized to the maximal peak-to-peak amplitude of the potentiated VL M-wave (RMS × M _max^-1^_).

### Antagonist coactivation level

The level of antagonist coactivation of the BF muscle (Co-Act_BF_, %) was computed as the BF EMG activity during knee extensions (KE), normalized to the maximal BF EMG activity recorded during a maximal knee flexion (KF). To record this maximal BF RMS value, the participants were asked to twice perform 3-s maximal voluntary isometric contractions of the KF before the fatigue protocol. This measurement was performed at a 90°-knee angle. The best trial was used for subsequent analysis.

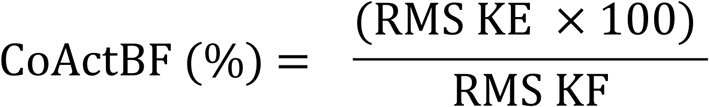

### Shear-Wave Elastography

An Aixplorer ultrasound scanner (version 12.2; Supersonic Imagine, Aix-en-Provence, France) coupled with a linear transducer array (4–15 MHz, SuperLinear 15-4; Vermon, Tours, France) was used in shear-wave elastography mode (musculoskeletal preset), as previously described [25]. Briefly, The SWE technique is based on ultrafast ultrasound sequences that are performed to capture shear-wave propagation. The SWE technique relies on the acoustic radiation force to remotely generate low-frequency shear-waves in tissues, (e.g., muscle, breast, liver), and can be achieved using the same piezoelectric arrays as those used in conventional ultrasonic scanners [25]. The shear-wave displacement field is saved via one-dimensional cross-correlation of consecutive radio frequency signals along the ultrasound beam axis as a function of time. The shear-wave speed is then calculated in each pixel of the resulting image using a time-of-flight algorithm on the displacement movies. The shear-wave propagation velocity, typically a few meters per second in soft tissues, correlates directly with the muscle shear elastic modulus (µ) if the medium is assumed to be purely elastic, which is well accepted in muscle elastography studies [25, 26]. The µ was obtained as follows:

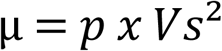

where *p* is the muscle density (1,000 kg*m^-3^) and V_s_ is the shear wave speed (in m*s^-1^). This equation implicitly neglects viscous effects. Studies revealed that the shear-wave velocity is almost independent of the frequency of the mechanical shock when measured longitudinally by SWE, demonstrating no significant viscous effects [26].

An ultrasound probe was placed on the VL muscle at 50% of the distance between the major trochanter and the lateral border of the patella. This position corresponds to the maximal VL ACSA. Then, the probe was aligned in the muscle shortening direction. The VL muscle was chosen because it is more sensitive to changes in stiffness than the head of the quadriceps muscle (*rectus femoris*) after fatiguing from long-duration exercise [22]. Moreover, studies showing decreases in muscle stiffness by assessing aponeurosis or muscle tendon junction displacements after repeated isometric contractions were made exclusively in the VL muscle [2, 16] allowing future comparisons with the present study. Shear-wave elastography measurements were carefully standardized by fixing the ultrasound probe using a custom-made system placed over the skin and coating it with a water-soluble transmission gel (Aquasonic, Parker laboratory, Fairfield, NJ, USA) to improve acoustic coupling. The B-mode ultrasound was first set to determine the optimal probe location and maximize the alignment between the transducer and the direction of the muscle fascicles. Probe alignment was considered to be correct when VL muscle fascicles and aponeurosis could be delineated without interruption across the image. A probe orientated in parallel to the muscle fascicles provided the most reliable muscle elasticity measurements [43]. After probe positioning, a fixed-size square region of interest (ROI; ∼1.5 cm^2^), *i.e*. a region in which shear-wave propagation was analyzed within the muscle, was placed in the middle of the B-mode image below the superficial aponeurosis within the VL muscle. A 2D real-time color map of the shear elastic modulus was then obtained at 1 Hz with a spatial resolution of 1×1 mm (**Fig 2A**). Finally, a 5-s clip was performed when the 2D real-time color map was maximally homogeneous to avoid any influence of the previous muscle contraction. During scans, the knee joint was positioned at 90° (0° = knee fully extended) and participants were asked to stay as relaxed as possible. Passive KE torque and VL and BF EMG activity were continuously and carefully monitored to verify this relaxed position.

**Fig 2.**
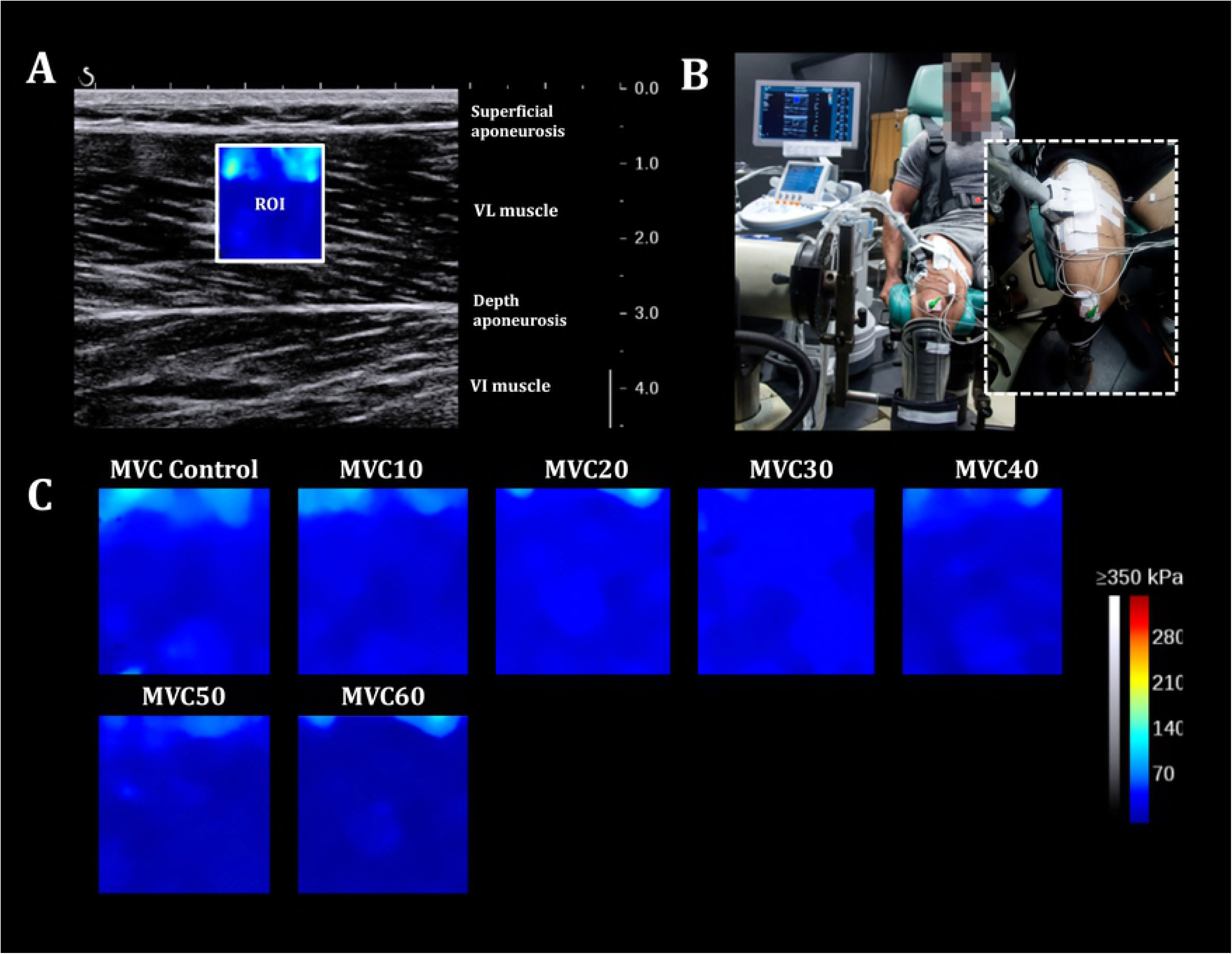
Shear-wave elastography. (**A**) 2D real-time color map of the shear elastic modulus in the *Vastus Lateralis* muscle obtained at 1 Hz with a spatial resolution of 1×1 mm. ROI: region of interest. (**B**) Participant positioning on the isokinetic ergometer with the probe fixation system placed over the right thigh. (**C**) A representative scheme of evolution of the 2D color map muscle shear elastic modulus for one individual.

The Aixplorer scanner provided Young’s modulus measurements, (E = 3 × *p* × V_s^2^_). The Young’s modulus calculation assumes that the material is isotropic, which is clearly not true for muscle [44]. Thus, all measurements were divided by three to obtain the shear elastic modulus (µ), as in the equation. The resting µ values were averaged over the largest ROI. The average of the five consecutive images (five values were recorded at one sample per second), obtained during the clip, was used for subsequent analyses. Reproducibility of the scans was also determined for each measurement. DICOM images were then transferred to a workstation and analyzed using a MATLAB ROI interface treatment script developed in our laboratory (MathWorks, Natick, MA).

### Statistical Analysis

The data were screened for normality of the distribution and homogeneity of variances using the Shapiro-Wilk normality and Levene tests, respectively. Differences in absolute values and relative changes (relative to control values) were analyzed by one-way ANOVA (effect: number of repetitions) with repeated measures. If the ANOVA revealed significant effects or interactions between factors, Fisher’s LSD *post-hoc* test was applied to test the discrimination between means. The intrasession repeatability of the resting shear elastic modulus was evaluated for the VL muscle between each five measurements, obtained during the 5-s movie, by calculating the intraclass correlation coefficient (ICC) and standard error of measurement (SEM) [45]. Results with a P value < 0.05 were considered to be significant. Statistical procedures were performed using Statistica 8.0 software (Statsoft Inc, USA). The results are presented as absolute values (mean ± SD) in **Table 1**. Data presented in **Figs 3-5** are expressed as the percentage of their initial value for the sake of clarity.

**Table 1.**
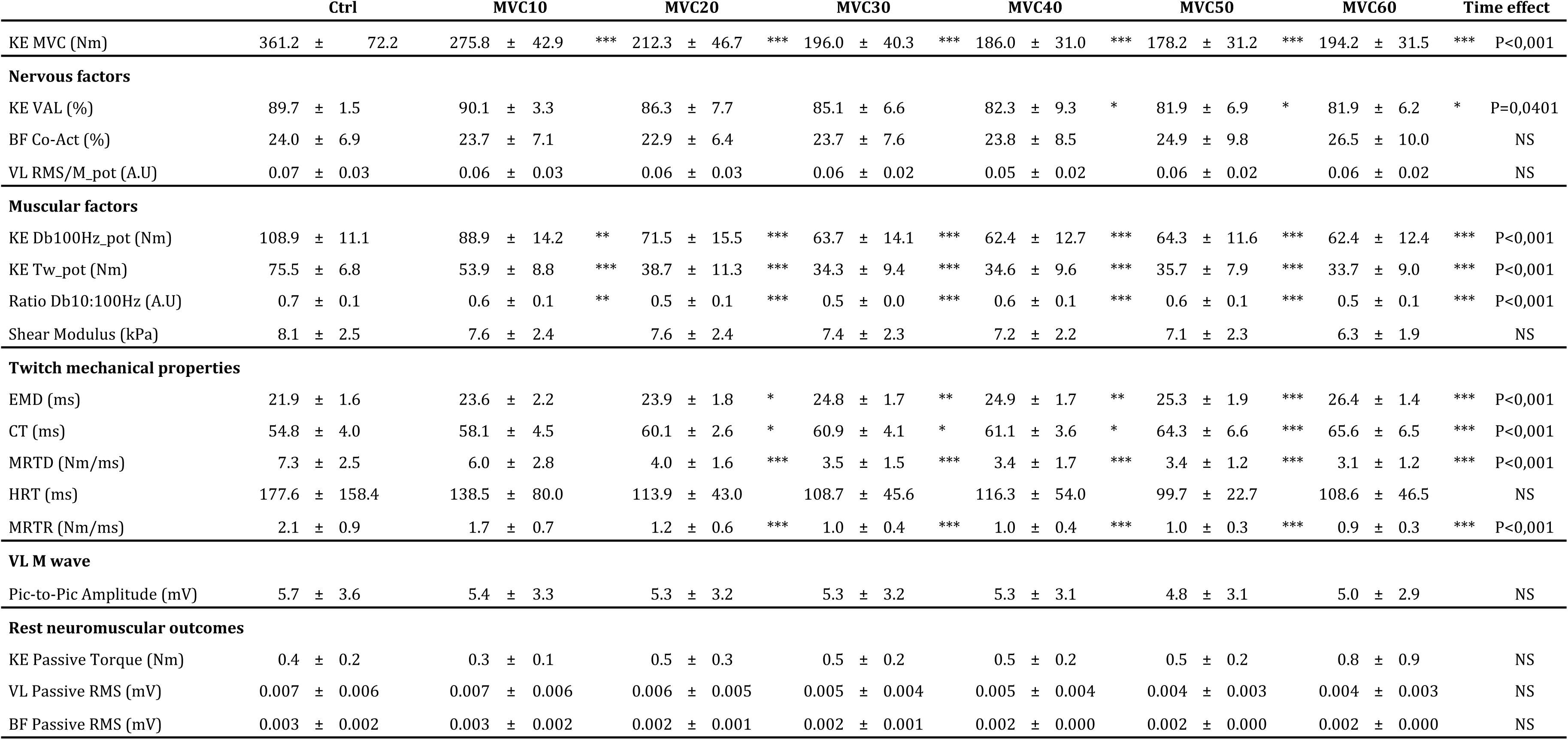
Kinetics of absolute neuromuscular and elastographic values. NS: Non significative result

**Fig 3.**
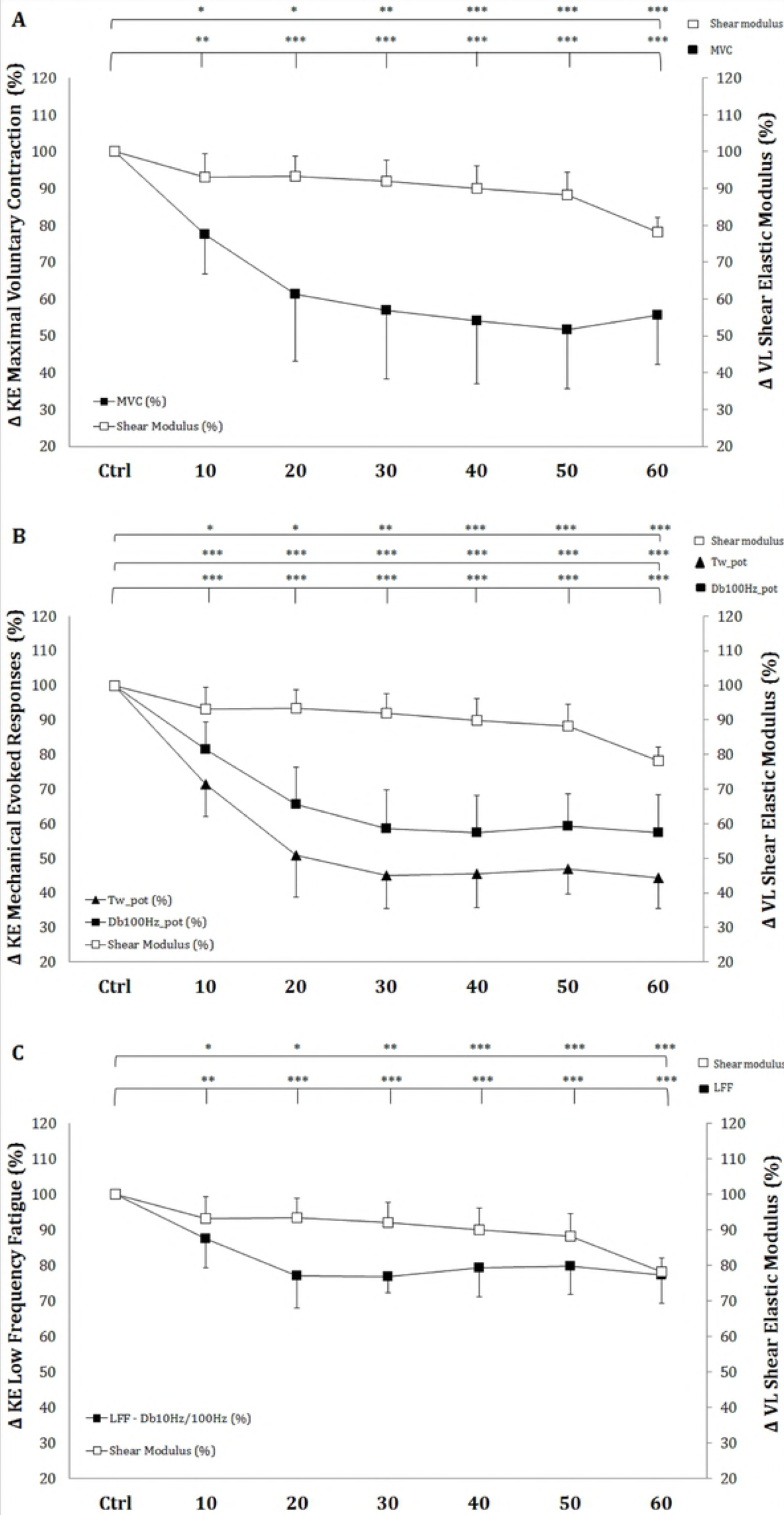
Time course of the voluntary and electrotimulated neuromuscular outcomes during the entire fatigue protocol. (**A**) Evolution (%) of the maximal voluntary contraction (MVC; ▪) amplitude. (**B**). Evolution (%) of double (Db100Hz__pot_; ▪) and single (T_w_pot_; ▴) pulses induced by electrical stimulation. (**C**) Evolution (%) of low-frequency fatigue (LFF; ▪), calculated as the ratio between the double pulses induced at 10 Hz and 100 Hz. The evolution (%) of the resting shear elastic modulus of the *Vastus Lateralis* (VL) muscle (□) is shown in each panel. Significant differences from the first MVC: * P < 0.05, **P < 0.01, and ***P < 0.001.

## Results

### Maximal voluntary torque

ANOVA revealed significant absolute and relative (to the control “non-fatigued” value) repetition-dependent effects on MVC torque (P < 0.001, **Fig 3A**). MVC decreased by 44.4 ± 13.4% by the end of the exercise (P < 0.001), with a significant reduction starting from the 10^th^ MVC (P = 0.0036).

### Peripheral mechanisms of fatigue

#### Electrically-stimulated potentiated torque

ANOVA revealed a significant repetition-dependent effect for both absolute and relative Db_pot100Hz_ and T_wpot_ torque similar to that of MVC (P < 0.001, **Fig 3B**). Db_pot100Hz_ and Tw_pot_ torque decreased by 42.5 ± 10.8% and 55.7 ± 8.8% by the end of the exercise, respectively (P < 0.001). Reductions in both relative Db_pot100Hz_ and T_wpot_ torque values started from the 10^th^ MVC (P < 0.001 for both), similarly to MVC.

#### Db10:100 ratio

ANOVA revealed a significant repetition-dependent effect for both the absolute and relative Db10:100 ratio (P < 0.001, **Fig 3C**). The Db10:100 ratio significantly decreased by 12.4% (P = 0.0012) from the 10^th^ MVC to 22.7 ± 8% (P < 0.001) by the end of the exercise.

#### Electro-Mechanical properties of single twitch torque

ANOVA revealed a significant repetition-dependent effect for both absolute and relative EMD and CT (P < 0.01, **Figs 4A** and **B**). The EMD and CT significantly increased during the fatigue protocol, reaching 121.1 ± 6.4% and 120.3 ± 14.5% of their initial values, respectively, (P < 0.001 for both) by the end of the exercise. Moreover, the values significantly increased from the10^th^ MVC for EMD (+8.0 ± 6.2%, P = 0.044) and the 20^th^ MVC for CT (+10.1 ± 7.7%, P = 0.0482). Moreover, the absolute and relative MRTD and MRTR significantly decreased by the end of the exercise: 56.7 ± 9% (P < 0.001) for the MRTD and 53.8 ± 14.7% (P<0.001) for the MRTR (ANOVA: P < 0.001 for both).

**Fig 4.**
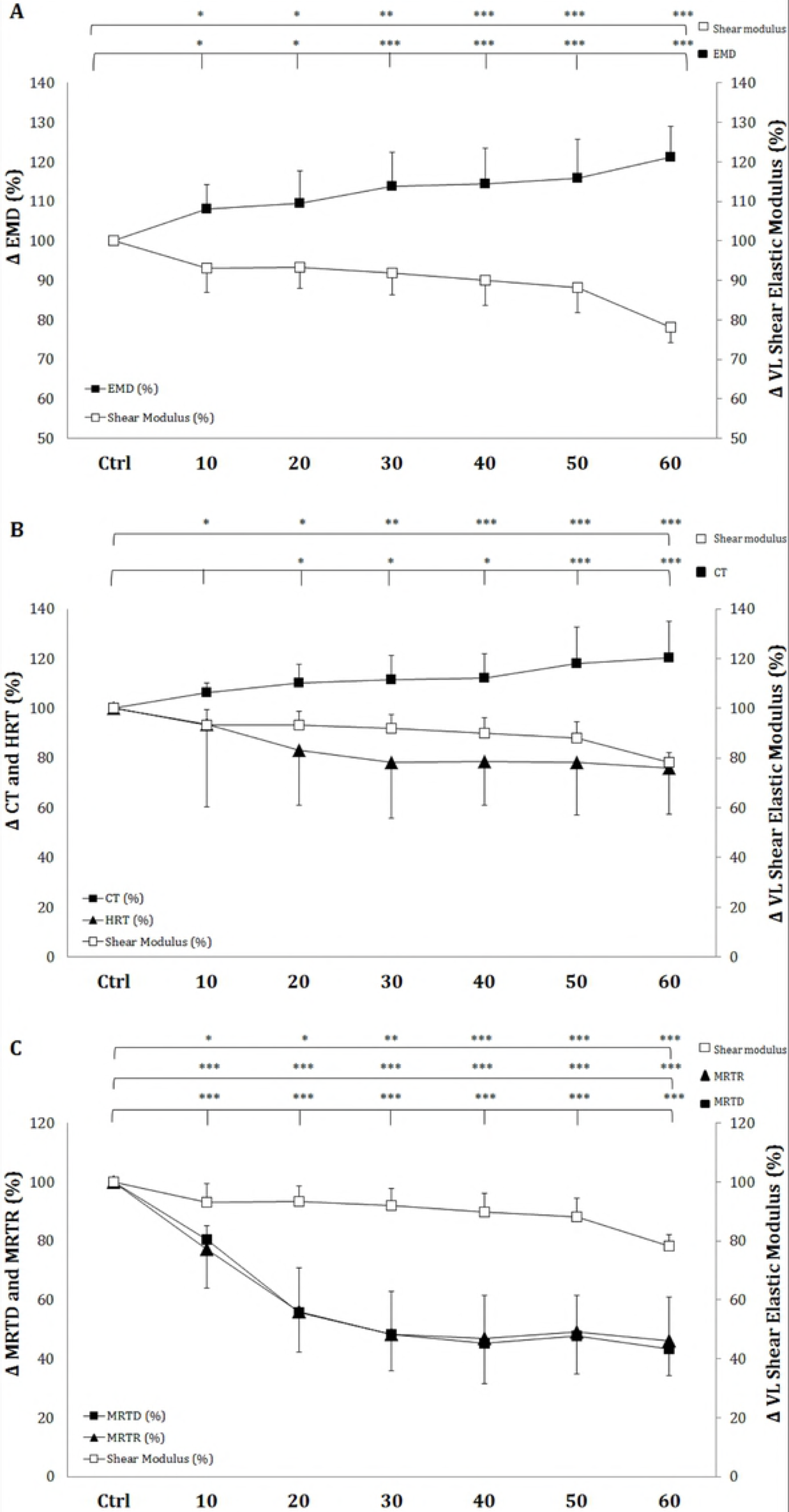
Time course of the electromechanical properties of the single twitch pulse during the entire fatigue protocol. (**A**) Evolution (%) of the electromechanical delay (EMD; ▪) amplitude. (**B**) Evolution (%) of the contraction time (CT; ▪) and half relaxation time (HRT; ▲). (**C**) Evolution (%) of the maximal rate of torque development (MRTD; ▪) and maximal rate of torque relaxation (MRTR; ▲). The evolution (%) of the resting shear elastic modulus of the *Vastus Lateralis* (VL) muscle (□) is shown in each panel. Significant differences from the first MVC: * P < 0.05, ** P < 0.01, and P < 0.001.

#### Potentiated Vastus Lateralis M-wave amplitude

There were no significant changes in the potentiated VL M_max_ amplitude during the entire fatigue protocol (**Table 1**).

#### VL resting shear elastic modulus

The measurements were reproducible (five measurements obtained during a 5-s clip) throughout the fatigue protocol. The ICC (95% CI) and SEM (kPa) values were as follows for each measurement: Control (99.5%, 0.2 kPa), MVC10 (99.5%, 0.24 kPa), MVC20 (98.9, 0.27 kPa), MVC30 (99.0%, 0.26 kPa), MVC40 (96.7%, 0.27 kPa), MVC50 (99.0%, 0.26 kPa), and MVC60 (97.5%, 0.26kPa).

The kinetics of the relative resting µ (in %) is presented in each figure. A representative scheme of the kinetics of the 2D color map of the muscle µ is presented in **Fig 2C**. ANOVA revealed a significant repetition-dependent effect for the relative resting VLµ (P < 0.001). It significantly decreased progressively from the 10^th^ MVC (-6.8 ± 6.2%, P = 0.0123) to the end of the exercise (-21.8 ± 3.9%, P < 0.001). The kinetics of the individual absolute values (kPa) of all subjects are presented in **Fig 5.**

**Fig 5.**
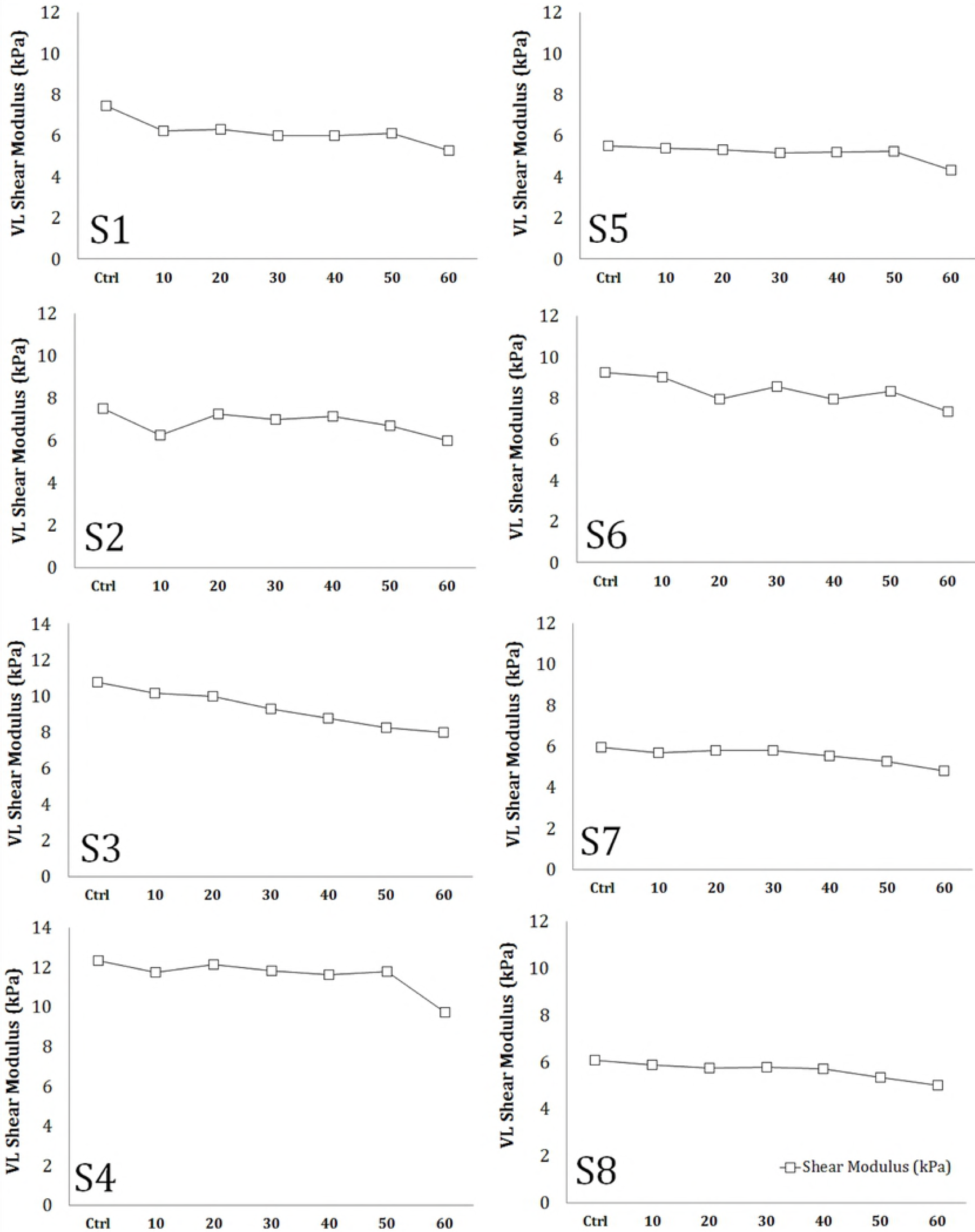
Time course of individual resting shear elastic modulus of the *Vastus Lateralis* muscle. The evolution of resting shear elastic modulus of the Vastus Lateralis (KPa) is represented for each participant before (Ctrl) and each 10 repetitions of the 10 x 6 sets of isometric maximal voluntary contraction of the knee extensor muscles of the fatigue protocol.

### Central mechanisms of fatigue

#### Voluntary activation level

Absolute VAL values are shown in **Table 1**. ANOVA showed a significant repetition-dependent effect for both absolute (P = 0.036) and relative (P = 0.015) VAL values. The KE VAL declined from the 40^th^ (-8.3 ± 9.8%, P = 0.016) to the 60^th^ MVC (-8.7 ± 8.8%, P = 0.011) at the end of the exercise.

#### EMG Activity

ANOVA revealed there to be no significant effect on the RMS·M _max^-1^_ ratio of the VL muscles nor the BF coactivation level over the entire fatigue protocol. Absolute values are presented in **Table 1**.

### Passive EMG activity and torque

Passive KE torque and VL and BF EMG activity assessed during the SWE measurements remained unchanged during the entire fatigue protocol **(Table 1).**

## Discussion

The aim of this study was to investigate whether the resting µ, evaluated by SWE, is a good indicator of peripheral fatigue during the repetition of isometric MVCs of the KE muscles in humans. We hypothesized that a series of isometric MVCs could induce a progressive decline in resting VLµ, reflecting greater muscle compliance and altered muscle function, and that modifications of resting VLµ may be directly associated with changes in electrically-stimulated mechanical (single or double pulses) and EMG (M-wave) responses. The main findings were that resting VLµ significantly decreased during the fatigue protocol from the 10^th^ MVC to the end of the exercise (60^th^ MVC) for all participants, with the loss ranging from -18 to -29%. The decrease in resting VLµ was also associated with decreases of voluntary torque, electrically-stimulated mechanical single and double responses, and the 10:100 Hz ratio (low-frequency fatigue). The VL M-wave amplitude was not significantly affected by the fatigue protocol. The kinetics of the resting VLµ were also associated with changes of the single twitch electromechanical properties. The EMD and CT increased significantly until the end of the exercise and the maximal rate of torque development/relaxation (MRTD and MRTR) were lower than control values. The HRT did not significantly change during the protocol.

### Reproducibility of measurements

SWE has been demonstrated to be a highly promising alternative to conventional elastography techniques, and provides a reliable and quantitative real-time assessment of muscular tissue stiffness at rest and during isometric contractions or passive stretching [21-23]. This study is the first to address the interest of µ on a non-contracted muscle at various stages of the fatigue during a series of maximal isometric contractions of the KE muscles. Methodological factors such as slight probe motion, compression, variability of the measurement, and the ability of subjects to achieve a fully relaxed state need to be meticulously controlled to ensure good reliability of the measurements [46, 47]. There was little variability of the measurements (five images obtained during the 5-s clip) during the entire fatigue protocol. ICC and SEM values varied from 96.7% to 99.5% and from 0.20 kPa to 0.26 kPa for the VL muscle, despite the development of fatigue. We achieved this level of reproducibility by positioning a custom-made probe fixation system over the skin and tightly taping it onto the VL muscle **(Fig 2B)**. In addition, the EMG VL and BF EMG activity and passive torque did not change during the SWE recordings **(Table 1)**, verifying that the participants were in a relaxed state.

Our resting VLµ baseline values are concordant with existing published SWE data [29]. The mean and standard deviation of 8.1 ± 2.5 kPa (ranging from 6.0 to 12.3 kPa) for a knee flexion of 90° (0° = knee fully extended) are in accordance with the values of < 10 kPa for the quadriceps muscles for the same knee flexion (90°) in active individuals in a non-fatigued state [29]. However, our control values were higher than those of other studies (∼3-5 kPa) for the VL muscle [22, 46, 48]. These differences may be explained by the short muscle length used in these studies (knee fully extended), as it is known that resting µ and muscle stiffness increase with increasing muscle length [27, 29].

### Muscle stiffness during and immediately after fatiguing exercise

An understanding of the characteristics of the muscle-tendon complex during and after fatiguing exercise has become essential for the prevention of over-use injuries [49]. Most previous studies have characterized fatigue or training-induced changes in muscle-tendon stiffness by exploring the movement of human tendons and/or aponeurosis structures *in vivo* [18-20]. Several studies have shown greater muscle-tendon compliance (increases in the elongation of connective structures for the same level of the produced force) after repeated contractions of the KE (e.g., VL muscle) and arm muscles [2, 16, 50]. However, one of the main limitations of these studies is that ultrasound-based techniques reflect modifications in the stiffness of several structures (muscles, tendons, nerves and skin) around a given joint and are not specific to skeletal muscle stiffness.

As mentioned above, SWE provides a quantitative and reliable measurement of individual muscular tissue stiffness [21-23]. A strong linear relationship between individual muscle force and dynamic muscle µ, evaluated during contractions, has been demonstrated, suggesting that it may be a good index of individual force [51]. Furthermore, Bouillard *et al.* [30] confirmed this relationship during submaximal isometric contractions, even when muscle fatigue occurs. These results suggest that SWE can be used to quantify relative modifications in voluntary force, even during fatiguing conditions. In another study [31], the same authors showed that when fatigue was previously induced in one quadriceps muscle (VL), lower dynamic µ values were observed, both initially and during a subsequent submaximal isometric task, relative to the control non-fatigued muscle. Although the amplitude of the muscle µ appears to be affected by fatigue, the underlying mechanisms for its decline are still unknown.

In our study, we observed a progressive decrease in the resting µ of the VL muscle from the 10^th^ MVC (-9.6 ± 8.5%) to the end of the fatiguing exercise (-21.8 ± 3.9%), suggesting a progressive rise in the VL muscle compliance. Some studies have suggested that increases in muscle-tendon compliance can be explained, in part, by alterations of the viscoelastic properties of the intramuscular connective tissue resulting from the increase in muscle temperature, due to repeated contractions [50, 52]. In our study, it is likely that decreases in the muscle µ amplitude were also accompanied by a rise in intramuscular temperature due to repeated maximal isometric contractions. Thus, modifications in the viscoelastic properties of exerted muscles due to fatigue may affect muscle performance. However, the kinetics of muscle temperature during and after the cessation of exercise and its association with the specific viscoelastic properties of muscles is still unknown. Currently, specific viscoelastic properties of the soft tissues may be quantified by real-time supersonic shear imagining, as described in the literature [25, 43], but no study has investigated the effects of peripheral fatigue on the viscoelastic properties of skeletal muscle and its role in force transmission.

The decline of the resting µ of the VL muscle found in our study is consistent with those observed for locomotor muscles after strenuous long-distance running, evaluated by invasive techniques, such as tension-myography and muscle belly deformation [24, 53], as well as SWE [22]. This exercise-model is clearly very different from that used in our study and the extent of induced fatigue may be greater than that in our model. For example, Andonian *et al*. [22] observed significant decreases in the resting µ of the quadriceps muscles (without distinguishing between the heads, but mainly in the VL muscle) after an extreme mountain ultra-marathon, which were still reduced after more than 45 hours. However, the relationship between changes in muscle µ and neuromuscular parameters of fatigue was not explored. In contrast to these results, Akagi *et al*. [54] observed an increase in resting µ (∼+7%) of the *medial gastrocnemius*, but not *soleus* or *lateral gastrocnemius,* muscle using another model of fatiguing exercise (1 h at 10% MVC). Lacourpaille *et al.* [29] showed that intense, non-damaging exercise (3 x 10 concentric MVCs at 120°*s^-1^) did not modify the resting µ of the elbow flexor muscles. They suggested that the resting µ would not be influenced by peripheral factors originating from fatiguing contractions, contrary to those observed in our study. However, in their study, 30 concentric MVCs did not induce a significant decrease in voluntary torque, limiting the conclusions concerning the relationship between fatigue and changes in resting µ. It is possible that in our study the significant decline in resting µ during the fatigue protocol may be closely related to the greater observed extent of exercise-induced peripheral fatigue. To date, there is no consensus among studies concerning modifications of muscle stiffness during and after fatiguing exercise. Methodological factors, such as the nature of exercise, duration, intensity, muscle length, morphological parameters of the subjects, and technical aspects, such as limb and probe positioning and the subject relaxation state could explain the discrepancies between studies.

### Shear elastic modulus and peripheral fatigue outcomes

The mechanisms underlying changes in muscle µ by SWE after fatiguing long-duration exercise are not well described [32]. However, several conclusions have been deduced from modifications in muscle stiffness after damaging exercise. Increases in muscle µ have been mainly observed minutes (30 min) and hours (48 h) after eccentric contractions [27, 29]. The authors hypothesized that increases in muscle µ may be strongly associated with the rapid perturbation of calcium homeostasis associated with this exercise modality. However, the relationship between the changes in muscle µ and cellular perturbations was not explored, limiting therefore conclusions. Moreover, no evaluations were made immediately after exercise in order to determine exercise-induced fatigue effects as propose in the present study.

We found that significant reductions in the KE MVC were mostly explained by alterations of peripheral rather than central factors (loss of -8.7 ± 8.8% VAL at the 60^th^ MVC). This is consistent with the literature, showing that high-intensity exercise induces greater peripheral [55] than central fatigue, which is mainly produced by long-duration exercise [40]. The large extent of peripheral fatigue observed was mostly reflected by a significant decrease in the amplitude of the single- and double-stimulated responses (-55.7 ± 8.8% and -42.5 ± 10.8%, respectively) from the 10^th^ MVC to the end of the exercise, but not by an alteration in the potentiated VL M-wave amplitude. Moreover, we observed low-frequency fatigue, *i.e.* the preferential loss of force at low frequencies of electrical stimulation, represented by a progressive decline in the 10:100 Hz ratio by 22.7 ± 8.0% by the end of the exercise. Modifications of this ratio are generally associated with changes in excitation-contraction coupling [1, 4, 8-11], more specifically, impairment of calcium homeostasis, *i.e.* decreased Ca^2+^ release, reuptake, and sensitivity. It has been hypothesized that increases in resting muscle µ following eccentric exercise can be attributed to alterations of calcium homeostasis due to structural disruptions [27, 29]. However, we propose that decreases in resting µ in response to fatiguing isometric contractions may be mainly associated with modifications of the elastic properties of the skeletal muscle responsible for force transmission and not by alterations of calcium homeostasis because of the exercise nature differences. The use of SWE at rest appears to be a quantitative and reliable measurement for exploring alterations of peripheral mechanisms during fatiguing contractions. However, the mechanism underlying decreases in resting µ under fatiguing conditions need to be investigated.

### Shear elastic modulus and force transmission properties

Single twitch properties, such as the EMD, CT, HRT, and their respective first derivates, the MRTD and MRTR, may also provide information concerning mechanical and electrochemical alterations of skeletal muscle related to force transmission in the fatigued state [12, 13]. For example, the EMD refers to the time between the onset of myoelectrical activity from the conduction of the action potentials along the sarcolemma to the development of tension originating from the contractile apparatus and stretching of the series of elastic components (electro-chemical and mechanical processes) [14]. EMD is considered to be a good indicator of the elastic properties of muscle, due to the important role played by elongation of the series of elastic components on total EMD [12, 56]. Indeed, it has been demonstrated that the EMD elongate by ∼15-20 ms after a fatiguing exercise of KE muscles (25 3-s isometric MVCs), and is associated with a temperature increase [14, 57]. These authors suggested that muscle compliance would increase due to muscular fatigue and the subsequent increase of muscle temperature, resulting in a longer time (a higher EMD) to stretch a more elastic muscle. In our study, the EMD significantly increased by 21.1 ± 6.4% (+ 4.5 ms) by the end of the exercise. Moreover, it was related to a significant increase in the CT (20.3 ± 14.5%) and decrease in the MRTD by 56.7 ± 9.0%. The MRTD inversely correlated with overall EMD after fatiguing exercise, in accordance with previous studies [14, 58]. Under fatiguing conditions, a lower MRTD would require more time to transfer tension to the tendon insertion point, thus increasing the EMD. In our study, we suggest that alterations on the electromechanical properties of single twitch may have been mainly associated with elastic rather than electrochemical processes (*e.g.* alteration in the propagation of action potentials along the muscle membrane), due to the lack of modification in the potentiated VL M-wave amplitude during the fatigue protocol. These findings strengthen the relationship between the decline in resting VLµ and alteration of muscle elastic properties (higher muscle compliance) with fatigue. Studies have suggested that fatigue-induced modification of the viscoelastic properties in the muscle-tendon complex could be strongly associated with a longer EMD (mechanical component) [15, 59]. These assumptions may explain the higher muscle compliance after fatiguing isometric contractions observed in this study. However, modification of the viscoelastic properties of skeletal muscle due to fatigue is yet to be investigated by supersonic imaging approaches. Thus, in the present study one head of the quadriceps muscle complex was evaluated limiting therefore interpretations about changes of the whole quadriceps neuromuscular properties with fatigue.

Finally, we also found a reduction of 53.8 ± 14.7% at the 60^th^ MVC in the relaxation phase after muscle contraction (MRTR) with the development of fatigue. It has been suggested that a more compliant elastic component in series requires more time to transmit the decline in cross-bridge tension to the tendon insertion point during relaxation, thus delaying the beginning of force decay [13]. Moreover, it could be related to the decrease in Ca^2+^ reuptake by the sarcoplasmic reticulum [60, 61]. Surprisingly, we did not observe any modification in the HRT. This suggests that alterations in the elastic properties of the skeletal muscle may have been mainly present during the development of tension originating from the contractile components and stretching of the elastic component in series rather than cross-bridge detachment. However, the high observed inter-individual variability in HRT values may explain this result.

## Conclusions

This study shows that the kinetics of resting VLµ may be associated with changes in both voluntary and electrostimulated torque responses and electromechanical properties of the single twitch during the repetition of maximal voluntary fatiguing exercise. Changes in resting VLµ may reflect a decline in muscle function, specifically the elastic properties, by increasing muscle compliance, directly affecting the capacity of force transmission. The use of SWE at rest appears to be a viable alternative and parallel tool to classical neuromuscular methods for the exploration of peripheral fatigue. However, the mechanism underlying the decrease in resting µ under fatiguing conditions still needs to be investigated.

This study provides scientific evidence related to changes of muscle µ associated with greater peripheral fatigue. However, it presents some methodological limitations: (i) the conclusions reported in the present study need be cautiously interpreted due to the small sample (n = 8). A more important sample is required in order to determine whether muscles stiffness is varying as a function of strength loss. Then, (ii) in the present study the whole quadriceps complex was not evaluated limiting interpretation of our results concerning others synergist muscles (RF and VM) and their association with the whole quadriceps neuromuscular assessments. Finally, (iii) it is possible that the magnitude of changes in muscle stiffness was different at other muscle lengths. Indeed, it has been suggested that muscle stiffness may vary as a function of muscle anatomical configuration (biarticular or monoarticular) [62] and muscle length (short or long) [27]. However, these findings were reported after eccentric but not fatiguing exercise. Thus, the relationship between muscle fatigue, stiffness and muscle length need therefore to be more investigated.

The SWE technique provides a non-invasive, quantitative, and reliable measurement of individual muscle tissue stiffness during fatiguing conditions. The study of both the muscle and tendon characteristics during and after fatiguing exercises in the context of sports medicine or the military is essential for the prevention of over-use injuries resulting from repeated exposure to low or high levels of force. More studies are required to confirm the observed higher specific muscle compliance with fatigue and its direct relationship with modifications of the stiffness of other non-contractile structures by ultrasound. Moreover, the *in vivo* analysis of the viscoelastic properties of skeletal muscle by supersonic imaging is necessary to better understand modifications in stiffness after fatiguing exercise and its impact on muscle performance.

## Acknowledgements

We thank Stéphane BAUGE, Stéphanie BOURDON, Philippe COLIN, and Benoit LEPETIT for their technical support and Pierre DEMAN for his contributions to SWE signal treatment. We also thank Oliver NESPOULOUS, MD. for his medical support.

